# Evolutionary implications of stipule occurrence in Permo-Carboniferous wetland forests dominated by Psaroniaceae

**DOI:** 10.1101/2025.06.23.661009

**Authors:** Weiming Zhou, Wenjun Sun, Ting Wang, Dandan Li, Josef Pšenička, Christopher Hill, C. Kevin Boyce, Jun Wang

## Abstract

Stipules are specialized appendages borne at the leaf base with roles as diverse as sheltering of delicate growing tissues from environmental exposure, vegetative propagation and dispersal, and modification into climbing hooks or protective spines. Stipules are widely present in extant vascular plants, but their origins in geological history remain obscure and must have involved convergent evolution more than once, perhaps as early as the late Paleozoic emergence of forested terrestrial ecosystems. Based on extraordinary collections from the early Permian Wuda Tuff Flora, the earliest known stipule structure in the plant kingdom has been identified in marattialean tree ferns Psaroniaceae, the dominant element of Permo-Carboniferous wetland forests. Psaroniaceous stipules are consistent with protection of juvenile fronds and the stem apex as well as retention of a functional role in mature and withered fronds. Furthermore, the continued and perhaps indeterminate growth and the fully laminated stipules invite speculation of a potential role in vegetative propagation following detachment from the parent frond. However, no direct fossil evidence of stipules acting as vegetative propagules is currently available.

## Introduction

The late Paleozoic was a pivotal period in the evolution of forested terrestrial ecosystems (DiMichele and Hook, 1992; Stein et al., 2012; Feng, 2017; Gastaldo et al., 2020). During the late Carboniferous, there was a shift in dominance from arborescent lycopsids (Lepidodendrales) to marattialean tree ferns (Psaroniaceae) in wetland forests (Phillips et al., 1974; Pfefferkorn and Thomson, 1982; Falcon-Lang, 2006; DiMichele, 2014). The high density of marattialean tree ferns with their broadly spreading canopy of fronds may have created some of the earliest closed-canopy forests (DiMichele and Hook, 1992), and also provided widespread microhabitats for small plants and animals (Rößler, 2000), thus playing a crucial role in the increasing complexity of forest ecosystems.

Stipules are specialized appendages borne at the petiole base. They are found widely in extant vascular plants and, at the least, can play a significant protective role for growing juvenile leaves along with the potential of a variety of more derived functions in specific lineages ranging from climbing hooks to vegetative propagation (Bell and Bryan, 2008). However, the origin and evolution of stipules have received little paleontological attention due to their rare fossil occurrence. In this study, based on the excellent preservation of whole-plant fossils, we confirm the presence of stipules in Psaroniaceae, overturning a long-term misunderstanding that these structures were absent (Stidd, 1971; Rothwell et al., 2018) since never previously found (Stidd and Phillips, 1968; Stidd, 1974; Hill and Camus, 1986). The form of these stipules at different developmental stages of psaroniaceous fronds suggests they may already have been involved in multiple functions.

## Results

From over 8000 fossil specimens of Wuda Tuff Flora, we found 149 specimens of aphlebiae consistent with interpretation as stipules, as explained below. While most specimens were preserved as isolated organs, 39 specimens show organic connection between aphlebiae and psaroniaceous trees. In a tree of *Scolecopteris libera* Li et al., fronds with intact aphlebiae at the petiole base constitute the outer periphery of the apical rosette (Figures 1A and B). A fallen psaroniaceous tree measuring 6.9 m was documented in a previous publication (Zhou et al., 2021). The lack of peripheral mature fronds makes the center of the stem apex more accessible, where aphlebiae are clearly present (Figures 1C and D).

**Figure 1.**
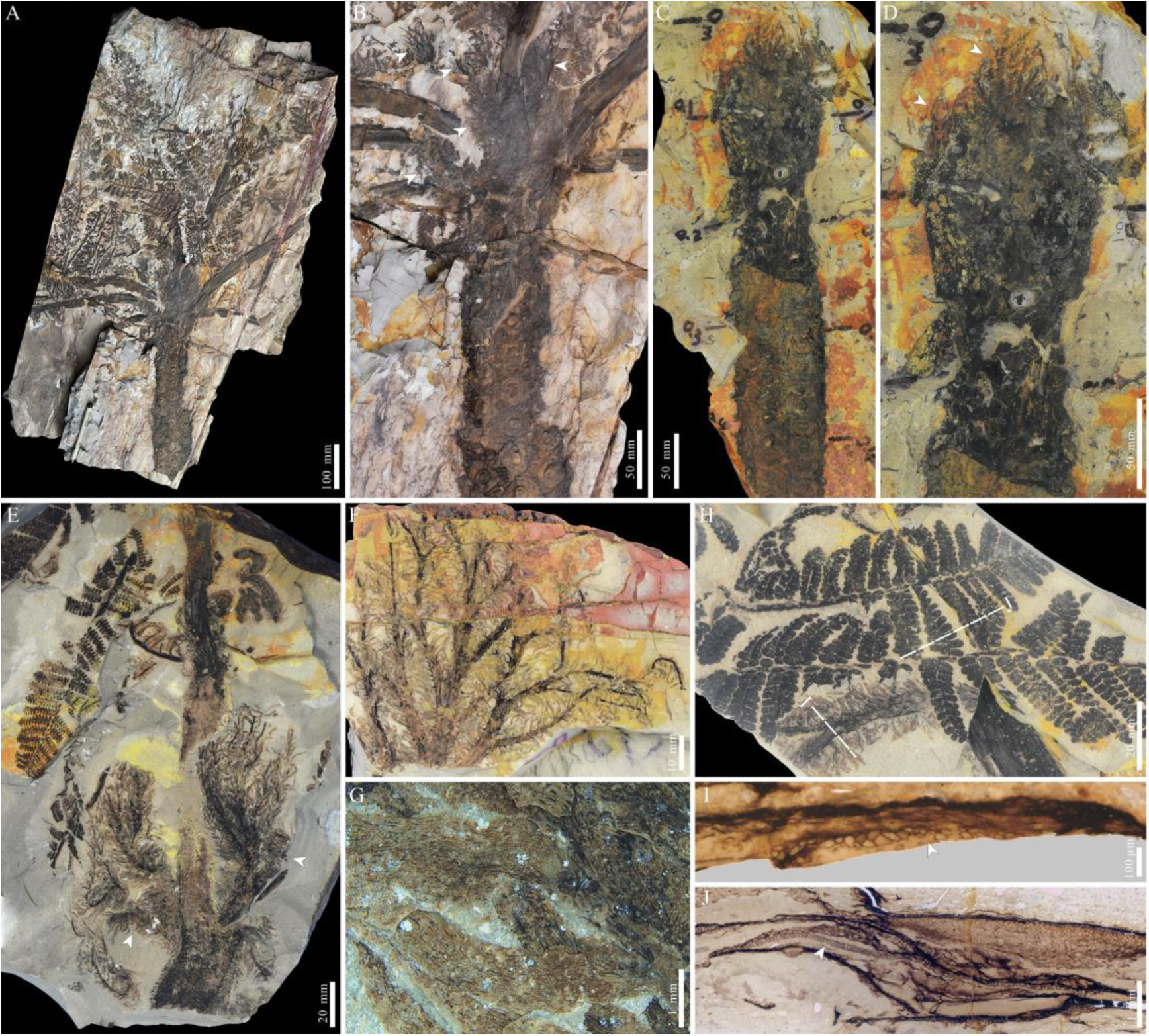
Non-laminated aphlebiae/stipules preserved isolatedly or in organic connection with psaroniaceous trees. (A) A psaroniaceous tree of *Scolecopteris libera*, specimen WH0002. (B) Enlargement of (A) showing aphlebiae attached to the peripheral fronds (white arrowheads). (C) A psaroniaceous tree lacking the preservation of peripheral fronds, specimen PB201856. (D) Enlargement of (C) showing the aphlebiae at the center of the stem apex (white arrowheads). (E) A psaroniaceous frond with paired aphlebiae at the petiole base, specimen PB201857. (F) Isolated aphlebiae with three orders of axes, specimen PB201858. (G) Aphlebia axis covered by non-vascularized scales, specimen PB201859. (H) A fragment of aphlebia associated with psaroniaceous fronds, specimen PB201860. (I) Anatomy of the aphlebia axis in (H), showing a transversely elongated xylem strand (arrowhead). (J) Anatomy of the associated psaroniaceous petiole in (H), showing a single main xylem strand (arrowhead).

Well-preserved specimens show that the aphlebiae were not grown directly on stems but appeared as paired structures positioned laterally on each side of the petiole (Figures 1E and 2). These aphlebiae are fan-shaped with three orders of branching (Figure 1F). None of the axes bear leaf-like laminae; instead, they are densely covered with tongue-shaped, veinless scales (Figure 1G). Anatomy of the second-order axis of an aphlebia reveals a single transversely elongated xylem strand (Figures 1H and I). Based on the macromorphology, the studied aphlebiae closely resemble the late Carboniferous species *Aphlebia hirsuta* (Lesquereux) White as both bear dense scales but lack laminae (White, 1899).

**Figure 2.**
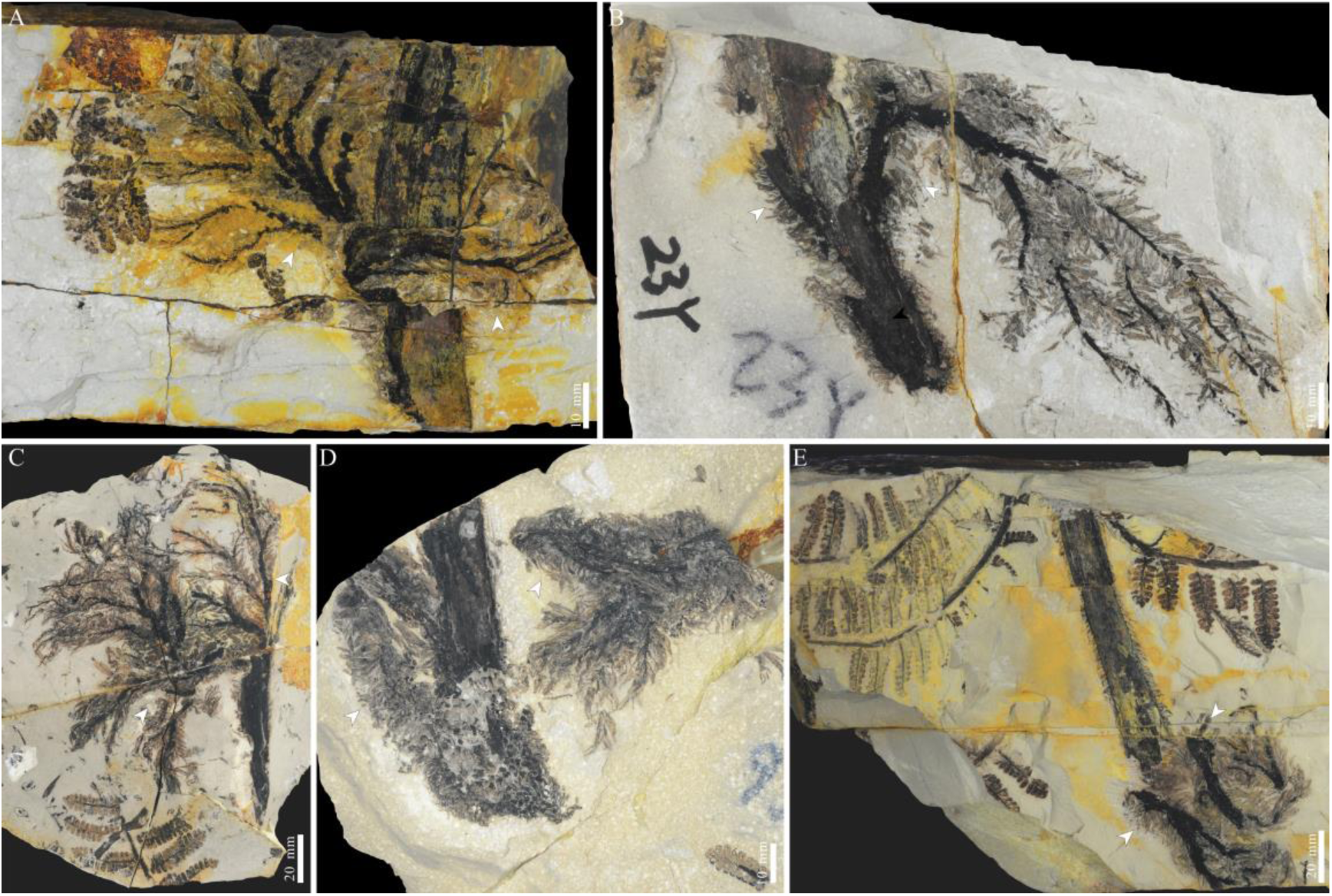
(A*–*E) Well-preserved specimens exhibiting non-laminated aphlebiae/stipules found in pairs at the base of petioles, specimen PB201864, PB201865, PB201866, PB201867, and PB204023 respectively.

In a psaroniaceous tree of *Pecopteris lativenosa* Halle, a new form of aphlebiae —slightly different from the aforementioned form by its thicker central axis and shorter lateral axes—is found at the petiole base of peripheral mature fronds (Figure 3A). Additionally, two block-like masses are observed at the center of the stem apex. Although their detailed morphology is indiscernible, some small circinate striations are visible in the left mass (Figure 3B). In another *P. lativenosa* individual, aphlebiae remain attached to the petiole base of a frond that appears withered, as inferred from its downward direction and its growth position well below the stem apex (Figures 3C and D). At the center of the stem apex, aphlebiae are apparently present, though their configuration is difficult to discern due to strong compression. On a local fracture surface, three larger circinate organs preserved across different bedding planes also exhibit internal circinate striations (Figure 3E). Through extensive examination, a coiled frond with similar internal striations is found in an isolated specimen, enclosed by a pair of aphlebiae with moderately developed laminae (Figures 3F and G). Notably, another specimen captures a later expansion stage in which vegetative growth of *Pecopteris*-type pinnules is evident (Figure 3H), confirming the affinity of this type of organ with Psaroniaceae. Taken together, these findings demonstrate that the larger circinate organs at the center of the stem apex in *P. lativenosa* (Figures 3B and E) are coiled fronds.

**Figure 3.**
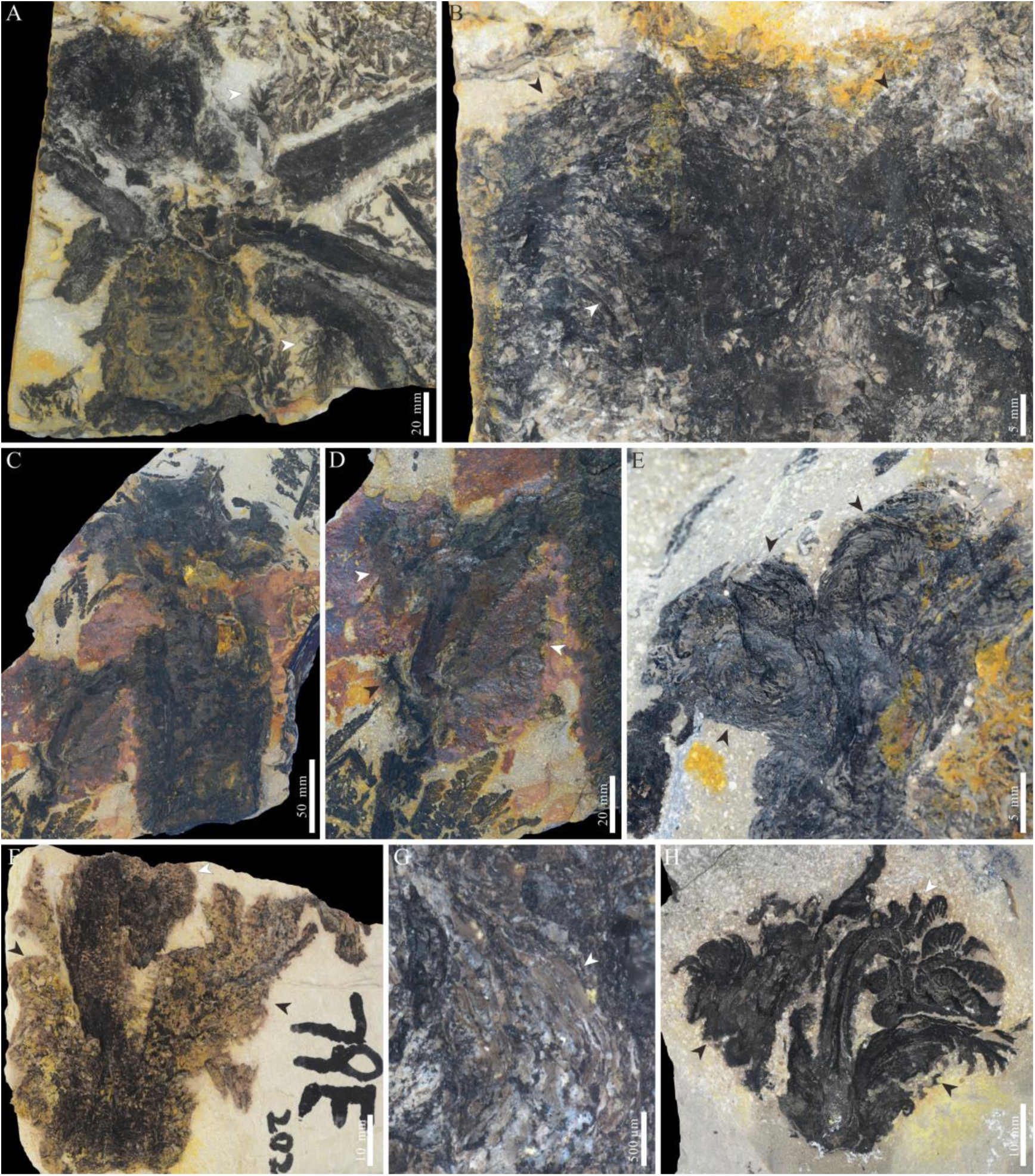
Coiled fronds preserved within psaroniaceous aphlebiae/stipules. (A) A psaroniaceous tree of *Pecopteris lativenosa*, showing a new type of aphlebiae (white arrowheads) attached to the petiole base of peripheral fronds, specimen PB203631. (B) Enlargement of (A) showing two block-like masses (black arrowheads), with the left mass exhibiting small circinate striations inside (white arrowhead). (C) Another psaroniaceous tree of *Pecopteris lativenosa*, specimen PB207381. (D) Enlargement of (C), showing a withered frond still attached to the stem. The paired aphlebiae are generally poorly preserved (white arrowheads), although the left one exhibits a clear apical morphology (black arrowhead). (E) Enlargement of (C), showing three large circinate organs preserved across different bedding planes (black arrowheads). Note the internal circinate striations. (F) An isolated specimen showing a coiled frond preserved within a pair of aphlebiae, specimen PB207382. (G) Enlargement of (F), showing small circinate striations within the coiled frond. (H) Another specimen showing a later stage of an uncoiling frond, still accompanied by a pair of aphlebiae, specimen PB207383.

## Discussion

### Identification of aphlebiae as stipules

The studied aphlebiae are distinct enough from other psaroniaceous organs as to have been mistaken during initial field identification for a lycopsid epiphyte occupying the *Psaronius* crown. However, we do not find these structures preserved in close association with any other trees of the forest community, as might be expected for an epiphyte. The key features of the studied aphlebiae with their vascularized axes and unvascularized leaf-like appendages (i.e., the scales) clearly distinguish them from the moss (unvascularized axes and leaves) and herbaceous lycopsids (vascularized axes and leaves) that have been documented in the late Paleozoic (Thomas, 1997; Hübers and Kerp, 2012). The studied fossils provide direct evidence of large aphlebiae that were borne near the base of psaroniaceous petioles a few centimeters distal from the point of abscission (Figures 1E and 2). The organic connection can be further supported by the fact that similar tongue-shaped scales densely cover the psaroniaceous petioles (Figures 1E and 2), and that the xylem strand of the aphlebia axis is comparable with that of associated psaroniaceous petioles (Figure 1J).

Despite being one of the best-known late Paleozoic plant groups (Taylor et al., 2009), Psaroniaceae failed to be recognized as stipulate due to several coincidences now evident in previous research. First, psaroniaceous stipules were previously found only as isolated organs identified as the fossil genus *Aphlebia* (Hill and Camus, 1986). Second, psaroniaceous stipules left no scars on the leaf cushions observed from the stem surface because they originated from the petiole rather than the stem (Blomquist, 1922). Third, psaroniaceous stipules were not recognized in some young sporophytes of *Psaronius* as this structure was absent in the first few leaves as the full size of the adult stem apex was progressively established (Blomquist, 1922; Stidd and Phillips, 1968). The present finding of psaroniaceous stipules highlights the advantage of whole-plant fossil preservation in the early Permian fossil-Lagerstätte of the Wuda Tuff, which can resolve centuries-old mysteries and controversies in palaeobotany (e.g., Wang et al., 2021).

The late Paleozoic tree fern family Psaroniaceae is widely accepted as the stem-group of Marattiales (Taylor et al., 2009). For comparison with the studied aphlebiae, *Angiopteris evecta* (Forster) Hoffmann is examined as a representative extant marattiaceous species. In *A. evecta*, paired stipules are borne a few centimeters distal to the petiole base (Figure 4A). These stipules are precociously developed, serving to protect the juvenile fronds during the later stages of its uncoiling process (Figures 4B and C). The vascular tissue of the stipule originates from the petiole and extends into the stipular lobes (Blomquist, 1922). As a consequence, transverse sections of stipules in more distal portions display multiple discrete vascular bundles similar to that of the attached petiole (Figure 4D). Scales are present on both stipules and petioles (Figure 4E and F). As paired, scaly, vascularized appendages of petiolar derivation borne in a similar position within the frond in two closely related groups, the aphlebiae of Psaroniaceae are interpreted as homologous to the stipules of Marattiaceae.

**Figure 4.**
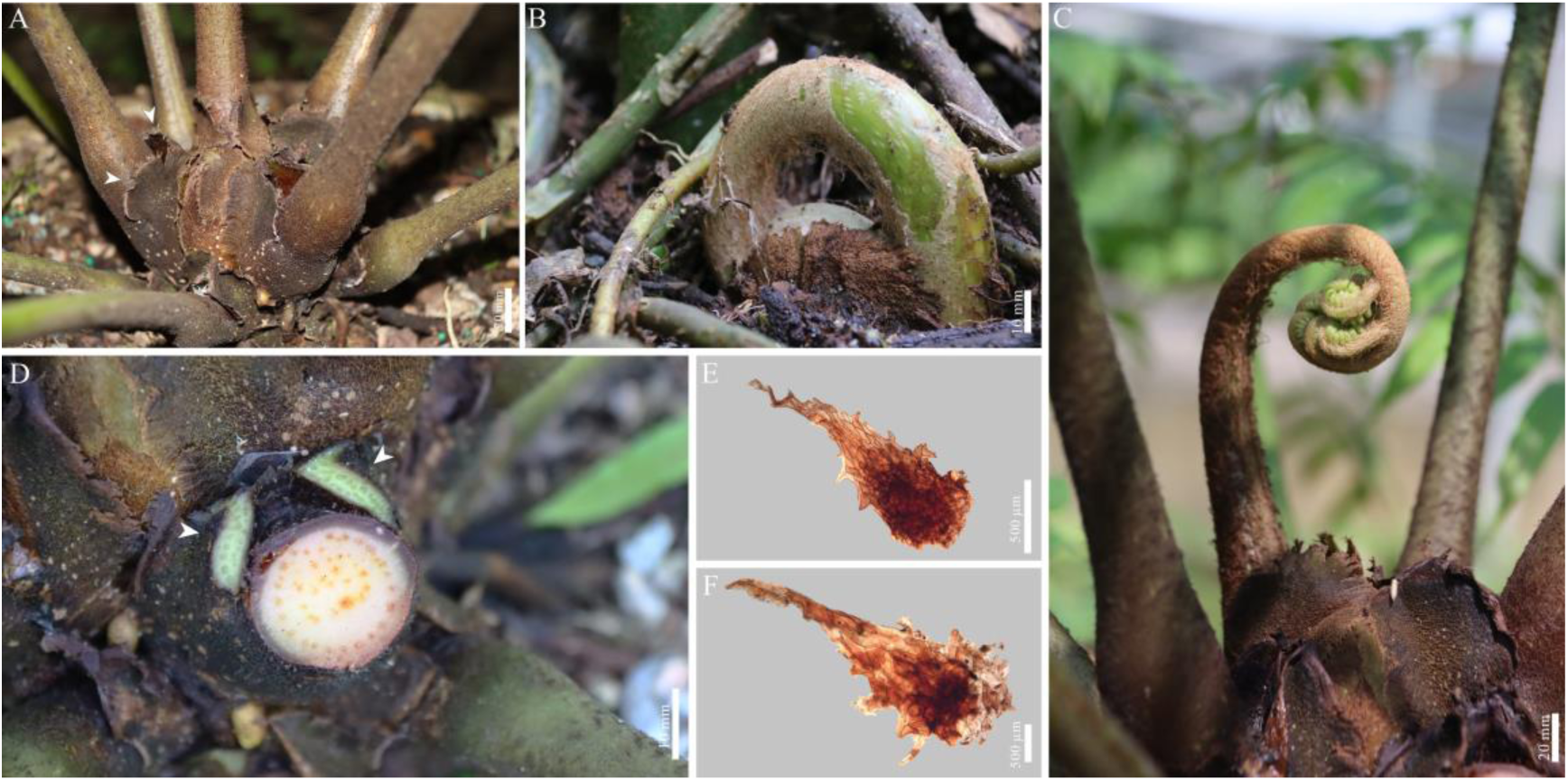
Stipule structures in extant *Angiopteris evecta*. (A) Paired stipules (white arrowheads) grown on the stem apex of *A. evecta*. (B) Coiled frond protected by paired stipules. (C) A later stage of uncoiling frond. (D) Anatomy of the frond petiole and attached paired stipules (white arrowheads), both with multiple discrete vascular bundles. (E*–*F) Scales collected from the connected portion of stipules and petioles.

The psaroniaceous stipules described here consisted of branching linear segments with a single vein per segment (Figure 1F). An evolutionary transition from this architecture to that of modern marattiaceous stipules involving multiple veins fanning out across a broad lamina (Figure 4D) would repeat a pattern seen across multiple convergences on a laminate leaf that reflect the developmental transitions between apical and marginal growth (Boyce and Knoll, 2002). For stipules to evolve laminate growth independently of the leafy pinnules on the same psaroniaceous fronds would be a striking instance of mosaic evolution, but not one without precedent, *e.g.*, modern *Ranunculus* staminodes evolved a laminate petaloid form independently of and developmentally distinct from the leaves of the same plant (Boyce, 2007). Notably, in this context, marattiaceous stipule anatomy in cross-section involves vascular bundles in a flattened ring, more like marattiaceous petioles than conforming to a single layer of bifacial bundles as in most leaf laminae (Figure 2B). The thick vascular bundle of psaroniaceous stipules is also more reminiscent of petiole vasculature (Figure 1K) than what is seen in the laminate pinnules (Stidd, 1974). In both Psaroniaceae and Marattiaceae endmembers, aphlebia/stipule anatomy would correspond to petiole anatomy, distinct from the anatomy and development of laminate pinnules.

While stipules are prevalent in contemporary angiosperms, a question arises regarding the timing of stipule origin, particularly in relation to the emergence of leaves. Late Paleozoic plant lineages that successfully developed laminate leaves included sphenopsids, ferns, progymnosperms, and seed plants (Boyce and Knoll, 2002). No evidence of stipules or analogous structures has been seen in the simple leaves of sphenopsids and progymnosperms. Among the frondose compound leaves of late Paleozoic ferns and seed plants, some have been identified to possess aphlebioid and cyclopteroid elements for which a function in the protection of young fronds has been postulated, similar to modern stipules (Bomfleur et al., 2012). However, these elements have been found also densely covering the stems and/or petioles (Phillips and Galtier, 2005, 2011; Krings et al., 2006). As such, they do not align with the strict definition of stipules and represent experimental developments of analogous structures. Psaroniaceae appears to be the earliest plant lineage to develop true stipule structures, long predating their widespread evolution in angiosperms. Given that leaves have evolved independently in ferns and seed plants (Boyce and Knoll, 2002; Tomescu, 2008), the stipule structures found in marattialeans and angiosperms must also have evolved independently.

### Stipule functions in psaroniaceous fronds

In extant Marattiaceae, stipules protect growing fronds, store nutrients, photosynthesize, and serve for vegetative propagation (Hill and Camus, 1986). Similarly, in psaroniaceous tree ferns, the precocious development of stipules early in frond ontogeny (Figure 1D), combined with the developmental trajectory of juvenile fronds as coiled forms (Figures 3B and 3E) expand and unfurl (Figures 3F and 3H), suggests that the dense aggregation of stipules at the center of the crown provided physical protection to the stem apex and juvenile fronds (Figure 5). In addition, the persistence of stipules in both mature (Figures 1B, 1E, 2, 3A) and withered fronds (Figure 3D) implies that they continued to serve functional roles throughout the frond’s lifespan (Figure 5). Furthermore, across the specimens studied here, psaroniaceous stipules exhibit a prolonged extension at the apex of segments that appears consistent with indeterminate distal growth (Figure 6A). This characteristic further distinguishes them from the aforementioned aphlebioid and cyclopteroid elements, which exhibit determinate growth. And the prospect of indeterminate growth raises the possibility of a role in vegetative propagation.

**Figure 5.**
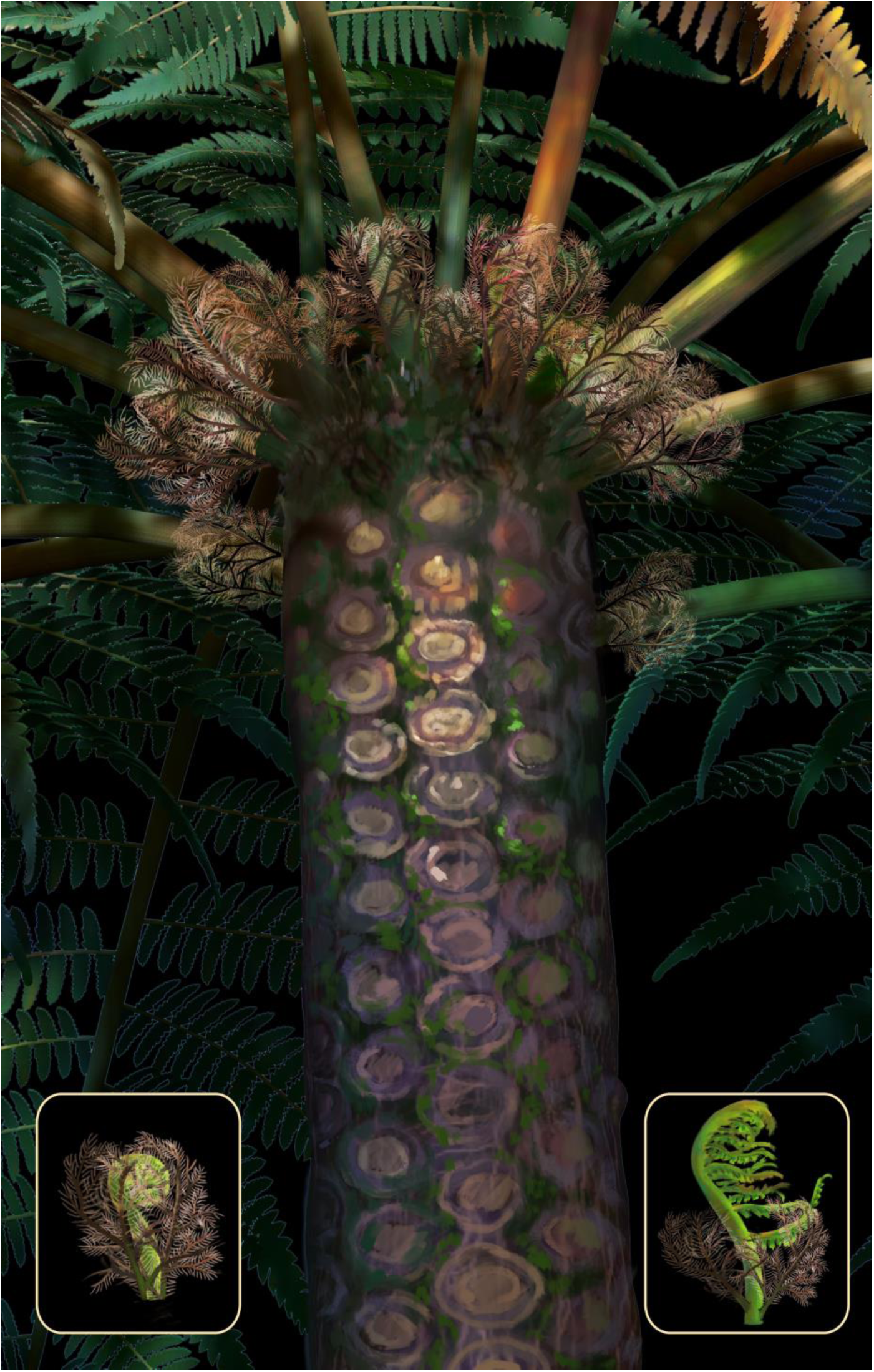
Reconstruction of aphlebia/stipule structures in late Paleozoic psaroniaceous trees. The lower-left box indicates a coiled frond preserved within the paired aphlebiae, while the lower-right box suggests a later stage with an uncoiling frond.

**Figure 6.**
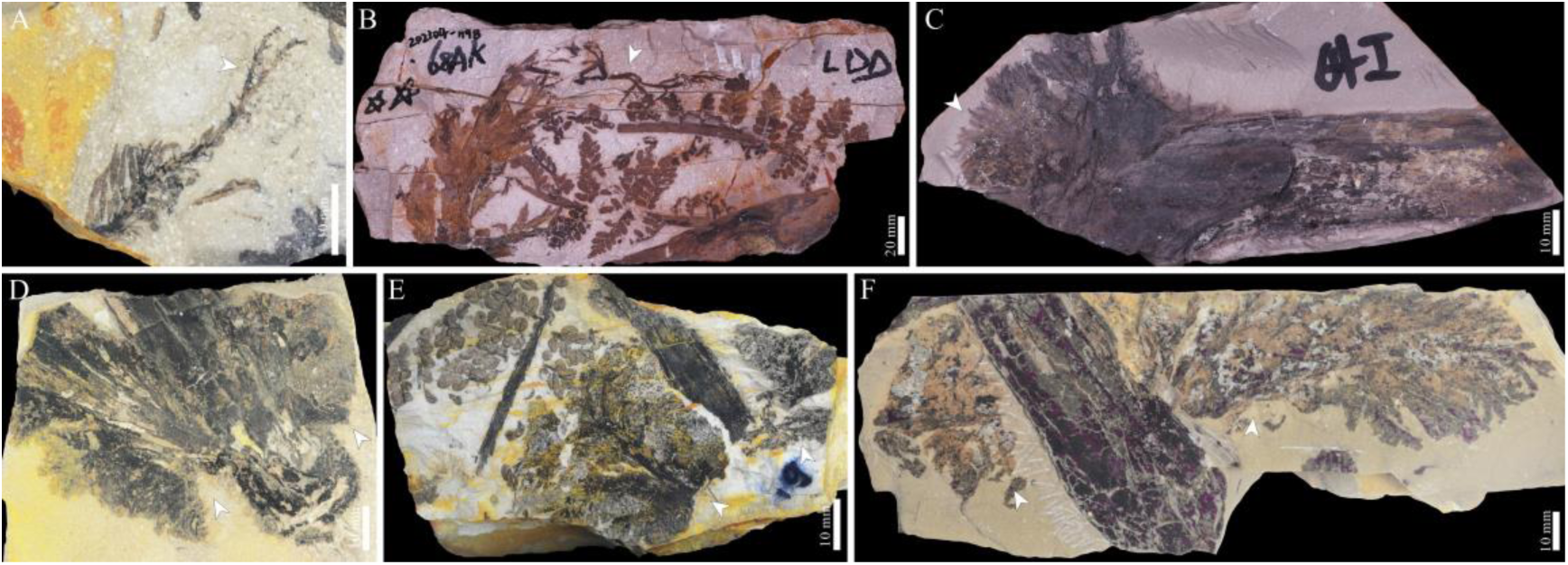
Variable lamination of aphlebiae, displaying characteristics akin to extant marattiaceous stipules. (A) A non-laminated aphlebia with continuous extension at the apex of the bifurcated segments (white arrowhead), specimen PB204024. (B) A moderately laminated aphlebia with continuous extension at the apex of the central axis (white arrowhead), specimen PB204025. (C) A fully laminated aphlebia, in round form with fringed margins (white arrowhead), specimen PB204026. (D–F) Moderately to fully laminated aphlebiae found in pairs at the petiole base (white arrowheads), specimen PB204027, PB204028, and PB204029 respectively.

The prospect of capturing the establishment for a psaroniaceous stipule as a vegetative propagule, complete with substrate interaction via adventitious roots, would be challenging. If such a process existed, it would only occur with detached fronds exposed on the substrate surface—an environment unfavorable for fossilization. Furthermore, psaroniaceous stipules do not appear likely to have been involved in photosynthesis at the preserved stages, as indicated by the absence of stomata in the epidermis of both axes and scales (Figure 1G). However, meristematic regions, representing rudiments of adventitious buds, have been observed in extant marattiaceous stipules (Blomquist, 1922) and vegetative propagation from stipules has been demonstrated experimentally to be more effective (Chiou et al., 2006; Huang et al., 2010) than the low rates of sexual propagation with poor spore germination and slow gametophyte growth seen in extant relatives (Chou et al., 2007).

Notably, several other types of aphlebiae with moderate to well-developed laminae have been found in isolation in the Wuda Tuff Flora (Figure 6) and other fossil localities in the literature (Hill and Camus, 1986). Although these aphlebiae are not preserved in a whole-plant context like the non-laminated psaroniaceous stipules studied here, Psaroniaceae should still be considered the most likely affinity of these structures for several reasons: 1) some aphlebiae display a similar pattern of extended apical growth (Figure 6B); 2) some aphlebiae that are round with fringed margins morphologically resemble the extant marattiaceous stipules (Figure 6C); 3) some aphlebiae occur in pairs at the petiole base (Figures 6D, E and F); 4) no other trees within this forest community are known to exhibit these aphlebiae. In particular, the presence of round aphlebiae (Figure 6C), which closely resemble the extant marattiaceous stipules (Figure 4A), further supports the idea that psaroniaceous stipules may have played a role in vegetative propagation.

Wetland environments can be home to seed plants with reproductive propagules that are unusually large (e.g. Paleozoic medullosans, DiMichele et al., 2006) or actively precocious (e.g. extant mangroves, Xu and Shi, 2024). Outside the lignophyte lineage that includes seed plants, Paleozoic wetlands saw the evolution of extreme heterospory in arborescent lycopsids and sphenopsids, and the only evolution of heterospory among ferns is the extant waterfern lineage. Large and/or precocious propagules may be essential given the stresses of establishment in a wetland environment with a saturated and anoxic substrate—as indicated by abundant organic matter preservation—that may at times have been fully submerged. No marattialean fern is known to have been heterosporous, but vegetative propagation via indeterminate growth in stipules as an asexual dispersal unit may have played a vital role in juvenile establishment and the overall success of the lineage in Paleozoic wetlands.

## Materials and Methods

Fossil specimens of Psaroniaceae were collected from the early Permian fossil-Lagerstätte Wuda Tuff Flora in the Wuhai City, Inner Mongolia, China. The Wuda Tuff Flora represents an early Permian swamp forest buried *in situ* by volcanic ash falls, allowing preservation of whole-plants and of community structure with high fidelity. Compositionally, psaroniaceous trees are often the dominant or subdominant group in different regions of the forest community (Wang et al., 2012). Fossil psaroniaceous trees are normally preserved as upright-standing stumps surrounded by frond litter but can also be found as horizontal tree falls possessing an intact distal rosette of fronds (Zhou et al., 2021). A spectacular example of the latter is specimen WH0002, now kept by the Suhaitu Coal Mine, Wuhai Energy, CHN Energy (Wuhai, China). Other specimens are maintained in the Nanjing Institute of Geology and Palynology, Chinese Academy of Sciences (Nanjing, China). Extant specimens of Marattiaceae were collected from the Orchid Conservation & Research Center of Shenzhen (Shenzhen, China).

The fossil specimen WH0002 was directly photographed in the field using a digital NIKON D7200 camera. Other fossil specimens were taken back into the lab, prepared with standard dégagement techniques, immersed in alcohol solution, and photographed using a digital NIKON D-800 camera under cross-polarized illumination. Two small pieces (with partial anatomy of psaroniaceous aphlebia and petiole respectively) were cut from specimen PB201860. Pieces were embedded in resin and prepared as petrographic thin sections using an EXAKT-300CP cutter and EXAKT-400CS grinder. Micromorphology of psaroniaceous slices and of aphlebia scales were observed and photographed using a Zeiss Discovery V16 stereomicroscope equipped with an Axiocam 512 digital camera. Photos of extant Marattiaceae were taken using a CANON EOS 90D camera. Scales of extant Marattiaceae were observed and photographed using a Nikon ECLIPSE E100 biological microscope equipped with a DFC 550 camera.

## Acknowledgments

This work was supported by the National Natural Science Foundation of China (42230210, 42172018), the Youth Innovation Promotion Association of Chinese Academy of Sciences (2022312), the Research Program of West Bohemian Museum in Pilsen (DKRVOZCM2020-25/92P), and the Visiting Professorship for Senior International Scientists of Chinese Academy of Sciences (2026PVA0102). We are grateful to Tang Jingjing, Yang Le and Dinghua Yang of Nanjing Institute of Geology and Palaeontology, Chinese Academy of Sciences for technical assistance.

